# Selection on cuticle color produces a genome-wide response that does not localize to known major genes in *Drosophila*

**DOI:** 10.64898/2026.09.29.755441

**Authors:** Sarah N. Ruckman, Anthony D. Long

## Abstract

Complex traits are often deemed highly polygenic when a mapping study fails to localize the loci underlying them, but a result that fails to identify major genes can also reflect a lack of power. Cuticle color in Drosophila offers a natural test of this distinction, since genes of large effect segregate in natural populations. We selected for dark and light thoracic trident color for 16 generations in two populations of *D. melanogaster* and two of *D. simulans*, then used pooled sequencing to scan each genome for allele-frequency differentiation between selected and control lines. Selection produced a strong, genome-wide shift in allele frequencies in every population, exceeding the expectation under drift. Simulations demonstrate that our design has low power to detect major-effect genes, which may explain why we do not observe peaks localizing to known color genes. We further show that a highly significant peak in each genome, on chromosome 3L, is an artifact of chorion-gene amplification in the pooled adult females. A source of false signal in pool-seq studies that has not been reported before. Cosmopolitan inversions shifted in frequency under selection in the one population that carried them, but direct testing showed they did not inflate genome-wide noise, contradicting a common assumption in the field. Together, these results demonstrate that a diffuse, polygenic-type response can arise from a trait with known large-effect genes, and provide a quantitative account of why an evolve-and-resequence design can fail to localize even a well-characterized genetic architecture.

## Introduction

A central challenge in modern biology is understanding the genetic architecture of complex traits. Such traits, including behavior, disease risk, and agricultural yield, arise from many interacting loci and from the environment (Ruckman and Long 2026). Identifying those loci is a shared goal of evolutionary biology, human genetics, and plant and animal breeding, and the methods for doing so have improved steadily, from classical quantitative genetics (Falconer and Mackay 1996) to linkage mapping in bi-parental crosses (Lander and Botstein 1989), mapping with multiparent populations, (King, Merkes, et al. 2012; King, Macdonald, et al. 2012), and genome-wide association (Visscher et al. 2017). Statistical power limits all of these methods. All genetic studies attempt to localize the genetic variance underlying a trait to specific positions in the genome. Most studies only account for a fraction of that variance, and the shortfall is termed missing heritability (Manolio et al. 2009). The standard explanation is that the remaining variance lies across many loci with individually small effects, each too small to meet a genome-wide significance threshold. Importantly, this explanation rests on what a study failed to detect rather than on positive evidence for the loci themselves. Complex traits are therefore routinely described as highly polygenic, or even omnigenic (Boyle et al. 2017; Visscher et al. 2017). A diffuse signal, however, can reflect detection limits as much as the trait’s biology.

Evolve-and-resequence (E&R) is a selection-based approach to the problem of localizing genetic variants affecting a trait (Schlötterer et al. 2015). In an E&R study, replicate populations are selected, pooled, and sequenced, and loci that respond show shifts in allele frequency relative to a control. Its appeal rests on a simple argument. Selection acts in every generation, so the change in allele frequency at a selected locus accumulates over the experiment. A locus of small effect shifts only slightly in a single generation, and that shift is hard to distinguish from the noise of sampling individuals and reads. Many generations of selection compound the shift, leading to a larger difference between the selected and control populations. E&R should therefore recover loci that a single-generation association study would miss. However, variance in allele frequency caused by drift also grows with each generation. At the small effective population sizes typical of laboratory selection, that variance grows quickly (Baldwin-Brown et al. 2014).

E&R has been applied to many traits in *Drosophila*, including accelerated development and lifespan (Burke et al. 2010), late-life fitness (Remolina et al. 2012), and adaptation to new laboratory and thermal environments (Orozco-terWengel et al. 2012; Tobler et al. 2014; Barghi et al. 2017). These studies typically return a broad, non-parallel response, spread across many loci that differ from one replicate to the next (Barghi et al. 2019). Such a response is generally read as evidence that the trait is polygenic (Barghi et al. 2020). However, low power produces the same pattern. The power of an E&R study depends on the number of replicate populations, the number of generations, the population size, and the linkage disequilibrium in the founding population (Baldwin-Brown et al. 2014; Kofler and Schlötterer 2014). The polygenic interpretation is rarely checked against the single-generation alternative that E&R is assumed to outperform.

Cuticle color may offer a way to separate power from architecture. Color is one of the few complex traits with known genes of large effect. In *Drosophila*, the melanin synthesis pathway is well characterized, and variants at genes in the pathway, including *yellow, ebony, tan, Adc, pale, burs,* and *bab*, alter cuticle color (Wittkopp and Beldade 2009; Takahashi 2013). The color of the thoracic trident, a dark marking on the dorsal thorax, is heritable and varies among wild populations (David et al. 1985). A pooled GWAS of natural variation also localized trident color to *ebony*, *tan*, and the cosmopolitan inversion In(3R)Payne (Endler et al. 2018). Color therefore could serve as a positive control for the ability of E&R to localize major genes. An adequately powered E&R scan should recover genes already known to be present. A diffuse result instead points to the power of the design rather than to the architecture of the trait, particularly when a single-generation design on the same trait recovers those genes.

We selected on trident color in two populations of *D. melanogaster* and two of *D. simulans* (Ruckman et al. 2026). For 16 generations, we selected the darkest and lightest individuals each generation to form dark and light selected lines, alongside an unselected control. Both species diverged in trident color. We then pooled and sequenced the dark, light, and control lines for each population at generation 16 and tested each SNP for differences in allele frequency among the pools, comparing the selected populations to the control. Most E&R studies concentrate their replicate populations within a single species. Instead, we divided our effort across two species, which allowed us to ask whether selection acts on the same genes in both species. Selection on a causal site also shifts the frequency of every site in linkage disequilibrium with it. A response therefore appears not as a single SNP but as a shifted haplotype block, often spanning hundreds of kilobases, and the causal site cannot be separated from the neighboring sites carried along with it. Populations of the same species share broadly similar haplotype structure, so a response at the same location in two *D. melanogaster* populations could still reflect a linked site rather than the causal one. *D. melanogaster* and *D. simulans* diverged roughly two to three million years ago (Obbard et al. 2012) and have recombined independently since. The two species share genes but not haplotype blocks. A response at the same gene in both species would therefore point to the gene itself rather than to a linked site. However, this design carries a cost. Power in an E&R study comes largely from replication, and dividing our effort across two species left two replicate populations per species, well below the number needed to localize selected loci (Baldwin-Brown et al. 2014; Kofler and Schlötterer 2014). We reasoned that color might be less affected by this limit than most traits, since its major genes have large effects and could rise above the background even with modest replication.

The known color genes did not stand out in our results. No single locus did. The response was broad and genome-wide, exceeding what drift alone would produce. This pattern matched the polygenic-looking response reported for other traits, even though past work has shown that large-effect color genes segregate in natural populations and can be uncovered in single generation case/control contrasts (Bastide et al. 2013; Endler et al. 2018). Modeling the power of our design against the noise in our own data explains why our intuition was incorrect. For a common allele, we had less than a 4% chance of detecting a major gene of any effect size. A single generation of case/control sampling is in fact a more powerful experimental design than a 16 generation selection experiment. We further show that the tallest peak in both species, on chromosome 3L, is an artifact of chorion-gene amplification in the pooled females rather than a response to selection. Chromosomal inversions are often blamed for E&R signals which fail to localize (Kapun et al. 2014), but they do not inflate the noise in our data, leaving replication and effective population size as the limiters on what we can reliably detect. Our results show that a diffuse E&R signal can arise in an experiment selecting on a trait segregating major genes.

## Methods

### Populations and Artificial Selection

The selection experiment is described in full in Ruckman et al. (2026). We selected on the cuticle coloration of the dorsal thorax in two species, *D. melanogaster* and *D. simulans*, each represented by two populations. We collected the *D. melanogaster* populations from Toronto, Canada, and Miami, Florida, USA, and refer to them by these localities. We collected both *D. simulans* populations from the same location in Tallahassee, Florida, USA, in different months of the same year, and refer to them as Tallahassee-1 and Tallahassee-2. The *D. simulans* populations are therefore temporal samples of a single locality rather than geographically distinct populations. At the start of the experiment, flies from each population were divided at random into three treatments: a dark selected line, a light selected line, and an unselected control. We then maintained each line separately for 16 generations, for a total of 12 lines.

Selection was gradual to limit bottlenecks early in the experiment (Kofler and Schlötterer 2014; Vlachos and Kofler 2019). For the dark and light lines, we scored the dorsal-thorax color of 250 females and 250 males each generation and bred the darkest or lightest individuals, breeding 80% of each sex in generation 1 and only the most extreme 8.4% by generation 16. Control lines were bred without regard to color, but experienced the same change in population size. The experiment ran for 16 generations, and as selection intensified, the number of breeding flies fell to roughly 42 by the final generation.

### Pooled sequencing and alignment

At generation 16, we pooled 350 females per line and extracted DNA from whole flies. Libraries were prepared with the Nextera kit and sequenced on an Illumina HiSeq as 150 bp paired-end reads. Reads were demultiplexed by barcode, aligned with BWA-MEM, sorted by coordinate, and assigned read groups with Picard AddOrReplaceReadGroups (pipeline at github.com/tdlong/XQTL2). For *D. melanogaster*, we aligned to the dm6 reference assembly (chromosome arms chr2L, chr2R, chr3L, chr3R, chrX). For *D. simulans*, we used the reference assembly 3.1 from NCBI. Since it carries NCBI chromosome names (for example, NC_052520.2), we renamed its chromosomes to the same chr-style scheme used in *D. melanogaster* (NC_052520.2 = chr2L, NC_052521.2 = chr2R, NC_052522.2 = chr3L, NC_052523.2 = chr3R, NC_052525.2 = chrX). All analyses used the five major chromosome arms.

### Coverage equalization by downsampling

Within each species, we analyzed each population independently across two pairwise contrasts: Light vs. Control and Dark vs. Control. Read depth differed among lines, and our test for differentiation scales with depth, such that lines sequenced more deeply show inflated differentiation given the same underlying allele frequencies. To control for this potential artifact, we downsampled the two lines in each population-by-contrast combination to the read count of the lower-coverage line. Downsampling was performed using samtools view with the subsampling option and a fixed seed of 42. Lines already at the minimum were retained unchanged. We then re-ran the entire variant-calling and testing pipeline on the downsampled data for each population separately.

### Variant calling and allele counts

We generated reference and alternate allele counts with the pipeline’s bam2bcf2REFALT step (bcftools v1.21). For each chromosome, we ran bcftools mpileup with indels skipped (-I), a maximum depth of 1000, and per-sample allele depths recorded, and piped the output to bcftools call in multiallelic-caller mode with variant sites only (-mv). We then retained biallelic SNPs with call quality QUAL > 59, removed any site with a missing genotype, and wrote per-sample reference and alternate read depths to a RefAlt table. The BAM list for each contrast contained only the downsampled experimental lines.

### Coverage analysis and chromosome X and 3L regions

We computed read depth in 1 kb windows for every line using samtools depth (counting all positions), recording the mean depth per window, and normalized to the genome-wide mean depth per line. We used this to evaluate whether genomic signals coincided with coverage anomalies. Two discrete regions showed coverage elevated several-fold above the genome average in all lines, across treatments, populations, and both species, each coincident with a cluster of chorion genes (see Results): a cluster on chromosome 3L (cytological 66D) and a cluster on the X (cytological 7F). Chorion genes are developmentally amplified in the ovarian follicle cells of females during oogenesis (Spradling and Mahowald 1980), and our pools consisted entirely of females, so the elevation in coverage reflects developmental gene amplification rather than a structural variant segregating among the populations. As a result, we treated both regions as amplification artifacts rather than targets of selection. We excluded them from the genome-wide inflation tests and displayed them in grey rather than removing them from the Manhattan plots, so that their coverage-driven signal is visible. The excluded windows were chr3L 8,500,000–9,000,000 and chrX 8,400,000–8,600,000 in *D. melanogaster,* and chr3L 8,342,753–8,834,090 and chrX 7,800,000–8,400,000 in *D. simulans*. All genome-wide claims rest on sites outside these windows.

### Tests of allele frequency differentiation

For each SNP, we tested for a difference in allele frequency between the selected line (dark or light) and the control line using Fisher’s exact test applied to a 2×2 table of reference and alternate read counts (fisher.test in R). We analyzed each population independently, yielding a genome-wide set of per-SNP tests for each population-by-contrast combination. Each contrast tested between 1.9 and 2.0 million SNPs in *D. melanogaster* and roughly 3.5 million in *D. simulans*, and slightly fewer after removing the chorion windows (1.91 to 1.99 million and 3.47 to 3.48 million, respectively). At each site, we recorded the p-value, odds ratio, and change in alternate-allele frequency between the two lines. We summarize genome-wide results as quantile-quantile plots of observed against expected -log_10_(*p*) and as Manhattan plots of -log_10_(*p*) against genomic position.

### Null simulations

To compare observed allele-frequency differentiation against theoretical expectations under drift and polygenic selection, we constructed null envelopes using a Wright-Fisher forward simulation, written in R (R Core Team 2025) with a fixed random seed (code at https://github.com/sruckman/Color_GWAS). We ran the simulation in 50 independent replicates under two conditions: drift only and variable polygenic selection. In both conditions, we clipped allele frequencies to [0.001, 0.999] each generation to prevent fixation.

In the drift-only condition, we initialized two independent populations with allele frequencies drawn uniformly from [0.05, 0.95]. Each generation, allele frequencies were updated via binomial sampling with 2*Ne* draws, where Ne followed the experimental schedule (Ne = 500 to 42 over 16 generations). We applied no selection to either line. The resulting p-values capture the expected allele-frequency divergence between two replicate lines under pure genetic drift with the same bottleneck history as the experiment.

For the polygenic selection condition, one line served as the unselected control and drifted as described above. The selected line received an additional per-generation change using the Falconer and Mackay (1996) truncating selection model: Δ*p*_i_ = *p*_i_(1 − *p*_i_) *i*(*t*) *s_i_*, where *i*(*t*) is the selection intensity at generation *t* (= φ(*x*_p_)/*p* where φ is the standard normal density and *x*_p_ = Φ⁻¹(1 − *p*)). We drew per-block selection coefficients from a Gamma distribution (shape = 2, mean s = 0.06) once per replicate. Population sizes and selection proportions per generation matched the experimental design (Ne = 500 to 42 over 16 generations). We converted simulated allele frequencies to read counts at 150× coverage and tested them with Fisher’s exact test, identical to the empirical pipeline.

We modeled LD among SNPs explicitly by grouping the simulated *n* SNPs into contiguous blocks of 100 SNPs each. We assigned each block a single selection coefficient drawn from the Gamma distribution described above, and updated the block’s allele frequency trajectory by selection each generation. Within a block, each SNP’s trajectory was a weighted average of the block-level trajectory and its own independent drift trajectory, with weights *w*(*d*) = exp(-*d*/40) and 1 - *w*, respectively, where *d* is the distance in SNPs from the block start. This gives pairwise LD ∝ exp(-*d*/40) within a block and approximately zero between blocks, where *d* is measured in SNPs. This short-range decay is consistent with the rapid breakdown of LD reported for natural populations of *D. melanogaster* (Langley et al. 2012). We drew starting allele frequencies for both block-level and per-SNP trajectories uniformly from [0.05, 0.95].

To build the envelope, we mapped each replicate’s sorted observed -log_10_(*p*) values onto a shared grid of 2,000 evenly spaced expected -log_10_(*p*) values spanning 0 to the theoretical maximum for *n* SNPs (-log_10_(1/*n*) ≈ 6 for *n* = 10⁶ SNPs per replicate). For each grid point, we estimated the corresponding observed value by linear interpolation between the two nearest data points. We then computed cross-replicate 5th, 50th, and 95th percentiles at each grid point to produce the ribbon and median line. We spaced grid points uniformly on the -log_10_(*p*) scale rather than the probability scale to ensure equal resolution across the full tail of the distribution.

### Power Model

To assess whether our design could detect a large-effect gene, we modeled the detection power for our experiment and a single-generation case-control design, following the framework of Long et al. (2026). The model contrasts the expected change in allele frequency at a selected locus (the signal) against the genome-wide variability in allele-frequency change (the noise).

The noise depends on a SNP’s starting allele frequency, so we estimated it as a function of that frequency. For each SNP, and separately for each contrast, we computed the change in alternate-allele frequency between the selected and control lines. We sorted SNPs by their starting frequency (i.e., their frequency in the control line) into intervals of width 0.02 and took the standard deviation of the frequency change within each interval. This standard deviation, pooled across chromosome-by-population strata and requiring at least 100 SNPs per interval, gives the noise as a function of starting frequency, σ(p_0_). We obtained uncertainty by resampling the chromosome-by-population strata with replacement (400 bootstrap replicates) and report 90% bootstrap intervals. The excluded chorion windows were omitted.

For the E&R design, we modeled the response to truncation selection over the 16 generations of the experiment. In each generation the allele frequency changed by i(t) *ε* p (1 − p), where e is the per-allele effect in phenotypic standard deviations, p is the allele frequency in that generation (starting from p_0_ and updated each generation), and i(t) is the selection intensity in generation t, calculated as i = φ(Φ⁻¹(1 − f)) / f for the proportion f of 500 scored flies bred in that generation (f = 0.800 in generation 1 to f = 0.084 in generation 16). The signal is the cumulative change after 16 generations. With R replicate populations, the noncentrality of the test scales as √R.

For the case/control design, we modeled a single generation of extreme-phenotype sampling. We assumed 1,000 flies, with the most extreme 5% pooled and sequenced at 100× or 200× coverage. The signal is i *ε* p_0_ (1 − p_0_) for a single round of selection, and the noise is √(p (1 − p) (1/N + 2/coverage)), combining individual sampling with read sampling.

We calculated power as the probability of exceeding a genome-wide significance threshold, using a two-sided normal test with a Bonferroni correction at α = 0.05. The correction used the number of SNPs tested per contrast in each species, roughly 2×10⁶ in *D. melanogaster* and 3.5×10⁶ tested in *D. simulans*. We evaluated power across per-allele effect sizes from 0.05 to 1.5 phenotypic standard deviations, at starting allele frequencies of 0.05, 0.15, and 0.35, and for R = 1, 2, 5, and 10 replicate populations. We modeled each species and contrast separately.

### Inversion frequency estimation

We estimated the frequency of seven cosmopolitan inversions in *D. melanogaster*, In(2L)t, In(2R)Ns, In(3L)P, In(3R)K, In(3R)Payne, In(3R)C, and In(3R)Mo, using inversion-diagnostic marker SNPs in dm6 coordinates from the marker set of (Kapun et al. 2014; Kapun et al. 2016). We ran samtools mpileup at the marker positions only against the dm6 reference, requiring base quality and mapping quality of at least 20 and suppressing indel and read-end output. A custom script counted the inversion-associated allele at each marker per line, and we estimated each inversion frequency as the mean of the per-marker frequencies. To test whether inversion frequencies changed during the experiment, we summed inversion-allele counts and total counts across all markers for a given inversion and line, then applied Fisher’s exact test to compare the control line with each selected line within each population. We restricted plotting and further analysis to inversions present at a frequency of at least 1% in at least one population in the control line: In(2L)t, In(3L)P, In(3R)K, and In(3R)Payne. This analysis was restricted to *D. melanogaster*; we did not estimate inversion frequencies in *D. simulans*, whose cosmopolitan inversion content differs.

### Inversions and Background Noise

We tested whether chromosomal inversions inflate genome-wide noise in allele frequency change, using two *D. melanogaster* populations. Miami segregates for In(2L)t, In(3L)P, and In(3R)K. Toronto does not carry these inversions. So, the two populations form an inversion-carrying vs. inversion-free comparison. We used the same noise estimate as in the power model: the standard deviation of allele frequency change as a function of starting frequency, σ(p_0_). We made two comparisons. First, between populations, we compared σ(p_0_) in Miami with σ(p_0_) in Toronto. Second, within Miami, we compared σ(p_0_) on the inverted autosomal arms (2L, 3L, and 3R) with σ(p_0_) on the non-inverted autosomal arm (2R). Toronto served as a negative control. It carries no segregating inversion on any arm, so any arm difference that appears in Toronto and Miami reflects a property of the chromosomes themselves rather than the inversions. We treated the X chromosome separately, since its reduced effective population size makes it drift faster than the autosomes, and excluded it from the arm comparison. We report σ(p_0_) at starting frequencies of 0.15 and 0.35. To estimate confidence intervals for the comparisons, we used a block bootstrap with 400 replicates and report 90% intervals and bootstrap p-values. For the between-population comparison, we resampled chromosome arms. For the within-population arm comparison, we resampled 500 kb windows within each arm. We excluded the chorion windows throughout.

### Population differentiation (Fst)

We measured genome-wide differentiation between the two populations within each species using Hudson’s Fst (Bhatia et al. 2013). We chose Hudson’s estimator because it is well suited to pooled sequencing. It is computed from allele counts, corrects for the finite read depth sampled in each pool, and is insensitive to unequal sample sizes between the two pools (Bhatia et al. 2013). We computed Fst between the two control lines, which represent the founding populations before selection: Toronto versus Miami in *D. melanogaster*, and Tallahassee-1 versus Tallahassee-2 in *D. simulans*. We used allele counts from the RefAlt tables and kept biallelic SNPs with at least 20 reads in each pool.

For each SNP, we computed the Hudson numerator and denominator, numerator = (p_1_ − p_2_)^2^ − p_1_(1 − p_1_)/(n_1_ − 1) − p_2_(1 − p_2_)/(n_2_ − 1) and denominator = p_1_(1 − p_2_) + p_2_(1 − p_1_), where p and n are the alternate-allele frequency and read depth in each pool. We averaged Fst across non-overlapping 10-kb windows using the ratio of summed numerators to summed denominators, with a minimum of 20 SNPs per window. We excluded the chorion windows, the same regions removed from the differentiation tests, and defined hotspots as the top 1% of windows by Fst within each comparison.

## Results

### Genome-wide differentiation exceeds drift

Selection produced a strong, genome-wide shift in allele frequencies. In both contrasts, Dark vs. Control and Light vs. Control, all four populations diverged far more than drift can explain (Figure 1). Every population sat above the drift-only envelope along the whole curve, not just at the tail (Figure 1). A polygenic selection simulation with a mean effect of s = 0.06 reproduced similar curves, with the populations falling near the envelope, some above and some below (Figure 1). The response was broad and genome-wide, coming from many sites rather than a few strong loci, consistent with the signature reported by other E&R studies of complex traits (Barghi et al. 2019; Barghi et al. 2020).

**Figure 1:**
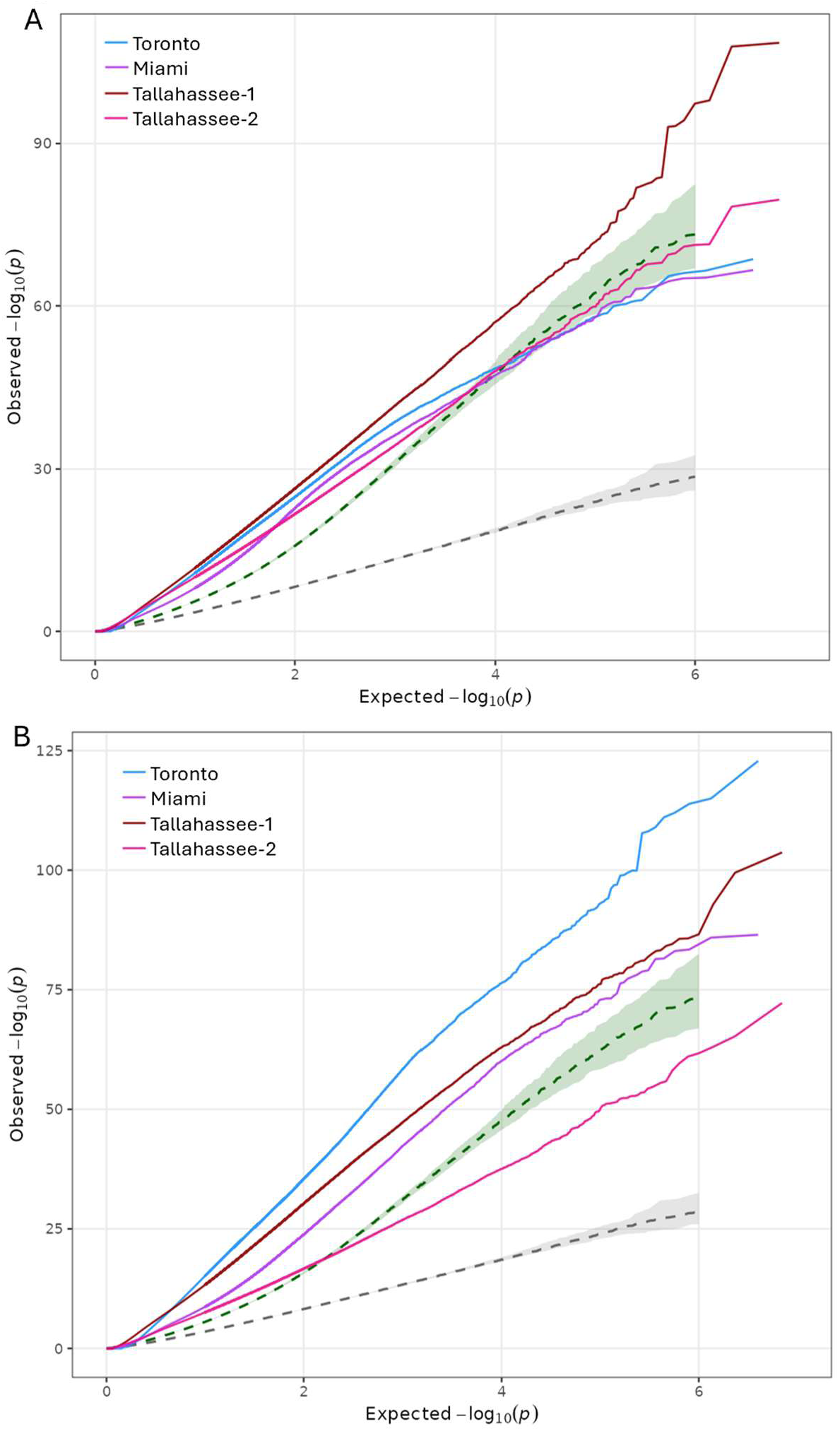
Quantile-quantile plots of observed against expected -log_10_(p) from Fisher’s exact tests of allele-frequency differentiation, for all four populations (Toronto and Miami, *D. melanogaster*; Tallahassee-1 and Tallahassee-2, *D. simulans*). The grey ribbon is the null envelope from a drift-only Wright-Fisher simulation matched to the experimental bottleneck (Ne = 500 to 42 over 16 generations); the green ribbon is a polygenic-selection simulation (mean per-locus effect s = 0.06). A) Dark vs. Control. B) Light vs. Control. Every population lies above the drift envelope across the whole distribution, indicating that selection shifted allele frequencies genome-wide.

### No single locus stands out

The response did not localize to any peaks. In either contrast, no single locus rose above the genome-wide background (Figures 2 and 3). The known large-effect color genes, such as *ebony* and *tan,* did not produce a peak in any population and were indistinguishable from the genomic background. The weak peaks that did appear were not shared among populations, and each population appeared to have its own peaks. However, there was one glaring exception: a tall, localized peak in all four populations on chromosome 3L, shown in grey in Figures 2 and 3.

**Figure 2:**
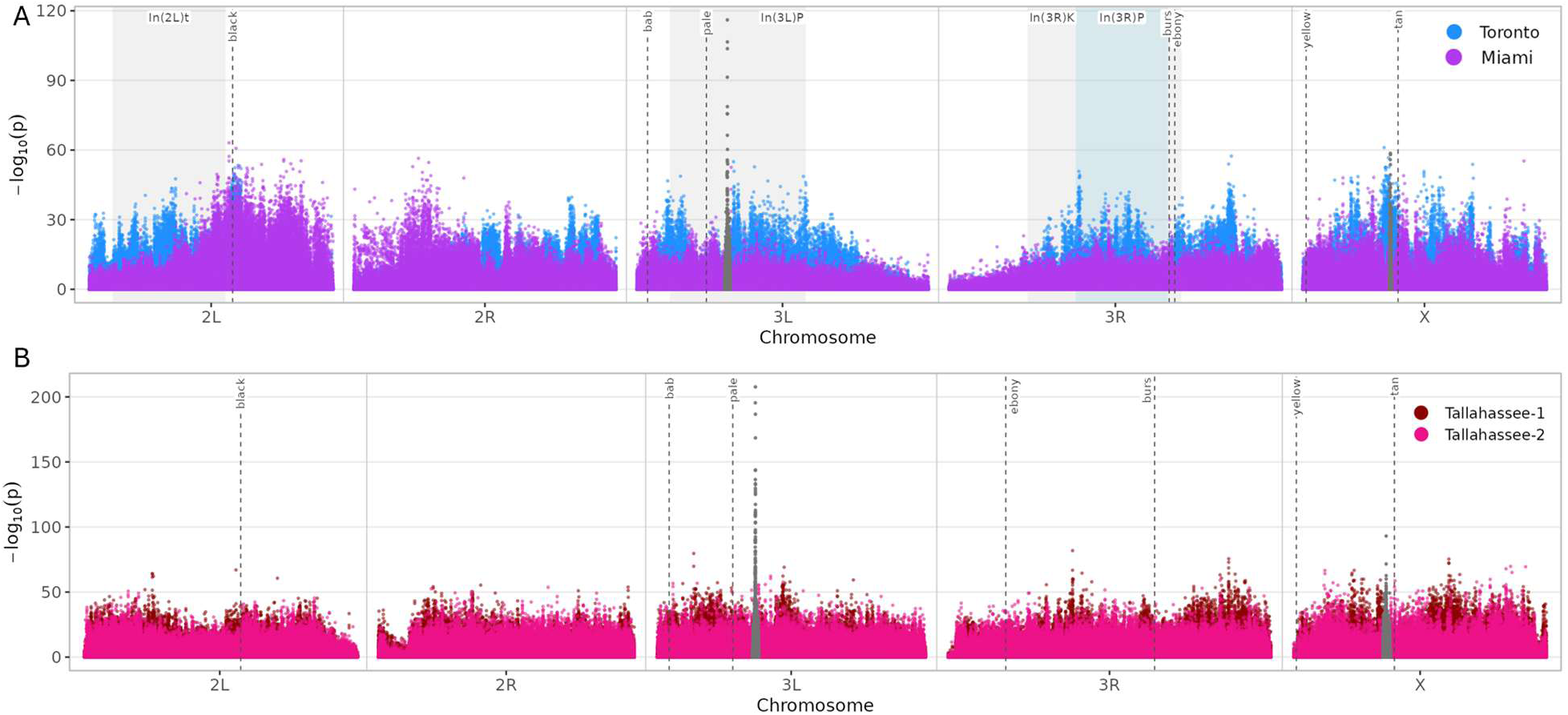
Manhattan Plots of Dark vs. Control. A) *D. melanogaster* B) *D. simulans*. Candidate color genes are marked with dashed lines, and cosmopolitan inversions are shaded in grey and blue (*D. melanogaster* only). The chromosome 3L and X chorion regions are shown in grey as coverage artifacts.

**Figure 3:**
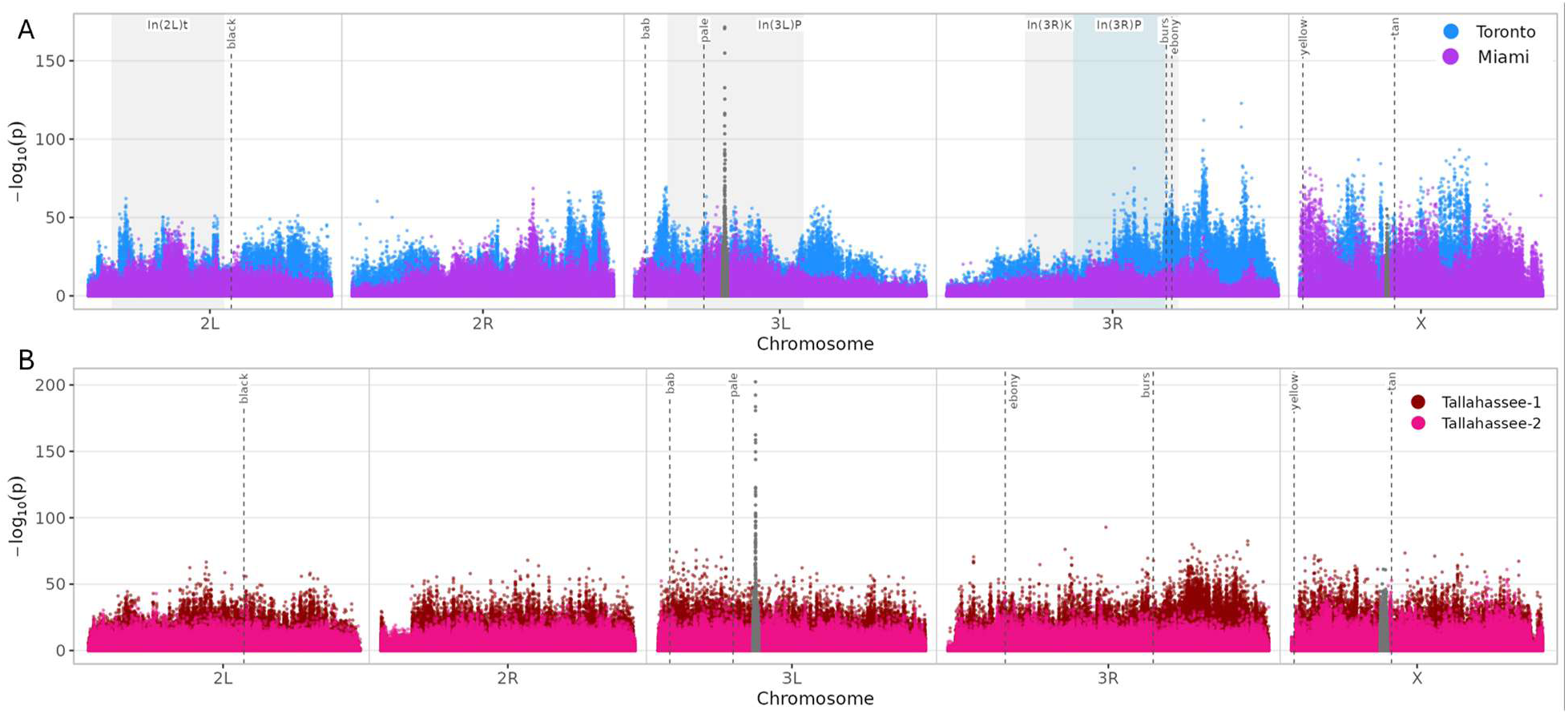
Manhattan Plots of Light vs. Control. A) *D. melanogaster* B) *D. simulans*. Candidate color genes are marked with dashed lines, and cosmopolitan inversions are shaded in grey and blue (*D. melanogaster* only). The chromosome 3L and X chorion regions are shown in grey as coverage artifacts.

### The major peak is likely due to chorion amplification in adult female DNA samples

The largest peak in the genome was not a response to selection. It sat on chromosome 3L in all four populations and both contrasts, over a cluster of chorion protein genes (chr3L 8.7–8.8 Mb in *D. melanogaster*; chr3L 8.5–8.6 Mb in *D. simulans*; Figures 4 and 5). Coverage was four- to five-fold above the genome mean across all 12 lines, the same in dark, light, and control, and in both species (Figures 4 and 5). The percent change in alternate-allele frequency at the peak was large and inconsistent. In *D. melanogaster*, it fell in Toronto (−80.7% light, −92.8% dark) and rose in Miami (+526.8% light, +133.0% dark). In *D. simulans,* it rose in Tallahassee-1 and Tallahassee-2 under dark selection (+7,230.4%; +48,533.3%) but fell under light (−91.1%) in Tallahassee-1. In Tallahassee-2 under light selection, the control carried no alternate-allele reads, so the percent change was undefined. Selection on a shared variant cannot drive changes in opposite directions across populations. Changes this large, and in one case undefined, cannot be read as a response to selection. Both mark the peak as an artifact. We therefore excluded the region from the genome-wide inflation tests (chr3L 8,500,000–9,000,000 in *D. melanogaster*; chr3L 8,342,753–8,834,090 in *D. simulans*) and show it in grey in Figures 2 and 3.

**Figure 4:**
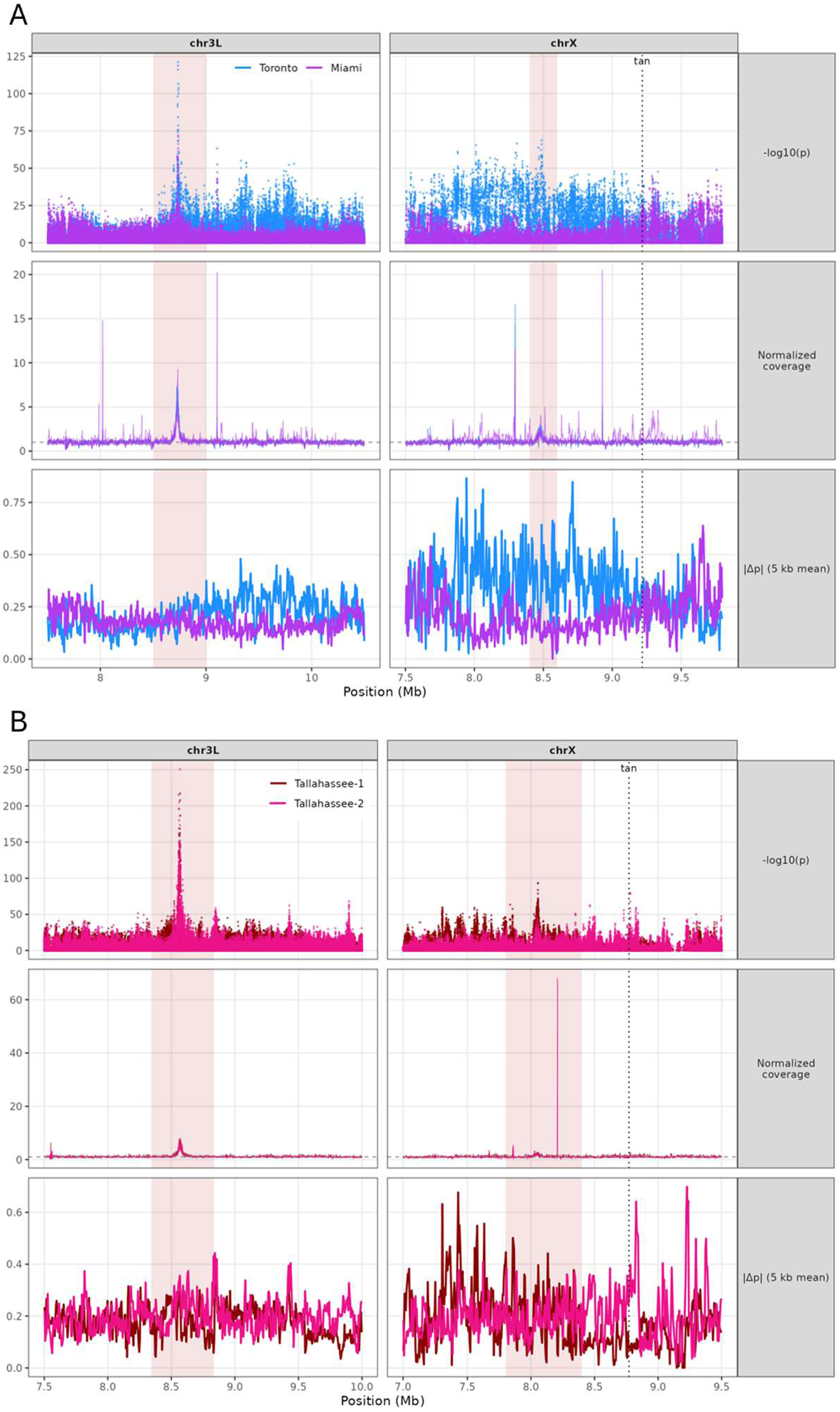
Coverage and association across the two chorion gene clusters, the 66D cluster on chromosome 3L and the 7F cluster on the X, in A) *D. melanogaster* and B) *D. simulans*, for the Dark vs. Control contrast. Rows: (top) −log_10_(p); (middle) normalized coverage; (bottom) absolute allele-frequency change in 5 kb means. Shaded bands mark the chorion clusters; the dotted line marks *tan*. Both clusters are elevated in both *D. melanogaster* populations, as predicted if the signal is developmental amplification in the pooled females, since a duplication on chromosome 3L would not predict elevation on the X. In *D. simulans,* the 3L cluster is clearly amplified in both populations, while the X cluster is elevated but variable.

**Figure 5:**
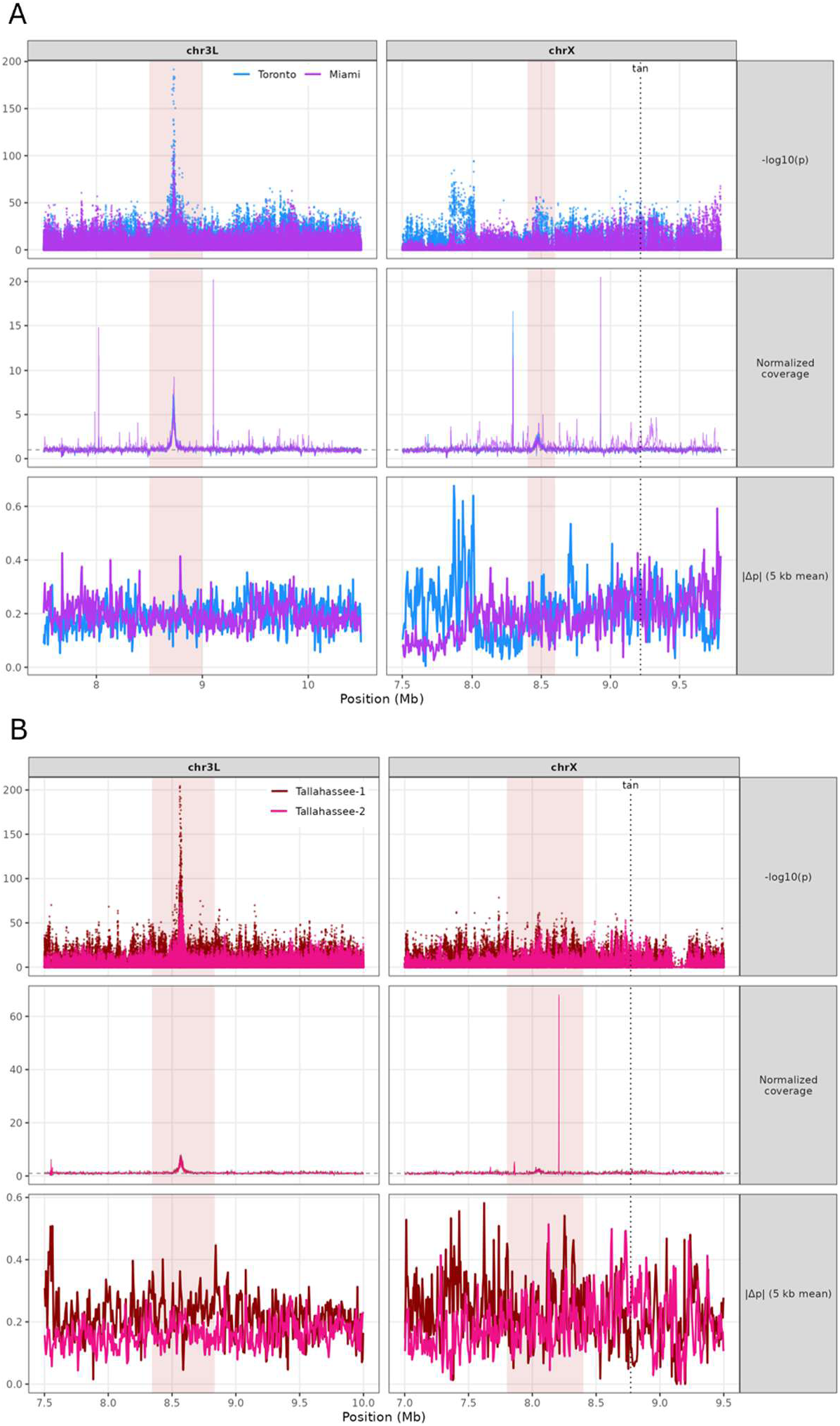
Coverage and association across the two chorion gene clusters, the 66D cluster on chromosome 3L and the 7F cluster on the X, in A) *D. melanogaster* and B) *D. simulans*, for the Light vs. Control contrast. Rows: (top) −log_10_(p); (middle) normalized coverage; (bottom) absolute allele-frequency change in 5 kb means. Shaded bands mark the chorion clusters; the dotted line marks *tan*. Both clusters are elevated in both *D. melanogaster* populations, as predicted if the signal is developmental amplification in the pooled females, since a duplication on chromosome 3L would not predict elevation on the X. In *D. simulans,* the 3L cluster is clearly amplified in both populations, while the X cluster is elevated but variable.

A direct test confirms that the elevation reflects developmental amplification rather than a segregating duplication. *Drosophila* carries two chorion gene clusters, and both are amplified in ovarian follicle cells during oogenesis: the third-chromosome cluster at cytological 66D, by roughly 60-fold, and an X-linked cluster at cytological 7F, by roughly 15-fold (Spradling and Mahowald 1980; Claycomb and Orr-Weaver 2005). Amplification in follicle cells therefore predicts that both clusters should be elevated in our all-female pools. A segregating duplication on chromosome 3L predicts nothing about the X. Both clusters were elevated in all six *D. melanogaster* lines, in dark, light, and control treatments alike. Peak coverage reached five- to nine-fold above the genome mean at the 66D cluster and two- to five-fold at the 7F cluster, in the rank order expected from their different levels of amplification in follicle cells. A segregating duplication on chromosome 3L cannot account for elevation at the X cluster, so the two-cluster pattern identifies the signal as follicle-cell amplification rather than a structural variant. The same pattern held in *D. simulans*, where both clusters were elevated in all six lines. The 66D cluster peaked at five- to eight-fold, and the X cluster was elevated in every line, though its magnitude varied widely (roughly four- to sixty-eight-fold). The two-cluster test therefore holds in both species. We also treated the X cluster as an artifact and excluded and greyed it alongside the 3L region (chrX 8,400,000-8,600,000 in *D. melanogaster*; chrX 7,800,000-8,400,000 in *D. simulans*; Figures 2 and 3). A duplication on chromosome 3L cannot explain elevation on the X, so the two-cluster pattern identifies the largest peak in each genome as chorion amplification rather than a response to selection.

### Our design had little power to detect a major gene

The E&R scan did not recover any known color gene, so we tested whether our design had the statistical power to detect one. We modeled detection power against the noise estimated from our own data, σ(p_0_), for our truncation-selection design at one to ten replicate populations and for a single-generation case/control design of 1,000 flies with the most extreme 5% sequenced at 100× or 200× coverage (Figures 6 and S1). We evaluated per-allele effect as the shift in mean phenotype in units of phenotypic standard deviation, produced by substituting one allele for the other, from 0.05 to 1.5 phenotypic standard deviations at starting frequencies of 0.05, 0.15, and 0.35 (Figures 6 and S1).

**Figure 6:**
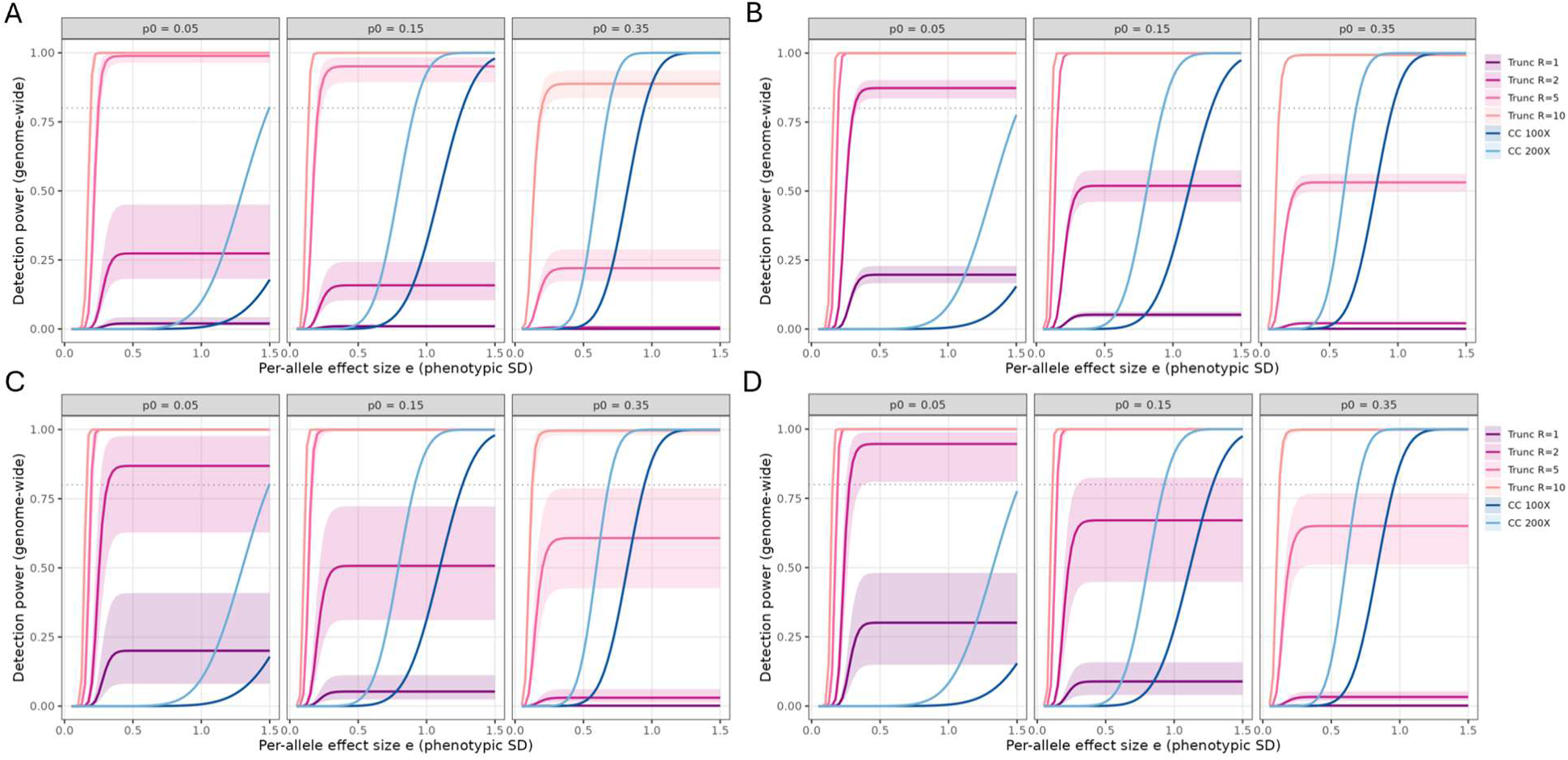
Genome-wide detection power (Bonferroni-corrected for 2 × 10⁶ SNPs) against per-allele effect size in phenotypic standard deviations, at three starting allele frequencies (p0 = 0.05, 0.15, 0.35), for A) *D. melanogaster* Dark vs. Control, B) *D. simulans* Dark vs. Control, C) *D. melanogaster* Light vs. Control, and D) *D. simulans* Light vs. Control. Curves show our truncation-selection design at R = 1, 2, 5, and 10 replicate populations and a single-generation case/control design at 100× and 200× coverage; bands are 90% block-bootstrap intervals over chromosome-by-population strata; the dashed line marks 80% power. E&R power saturates with effect size and, at our replication (R = 2), never exceeds about 4% for a common allele, whereas case/control power continues to rise.

In the E&R design at our replication level (two populations per species), detection power rose with effect size and then plateaued above a per-allele effect of roughly 0.4 phenotypic standard deviations, so larger effects added little power (Figure 6). Maximum power to detect a major gene remained low. For an allele at a starting frequency of 0.15, the maximum power across effect sizes was 0.16 in the Dark vs. Control contrast and 0.51 in the Light vs. Control contrast in *D. melanogaster*, and 0.52 in the Dark vs. Control contrast and 0.67 in the Light vs. Control contrast in *D. simulans*. For an allele at a starting frequency of 0.35, maximum power did not exceed 0.04 in any comparison (*D. melanogaster*: 0.006 Dark, 0.030 Light; *D. simulans*: 0.021 Dark, 0.033 Light).

Power in the single-generation case/control design rose with effect size without plateauing and reached 1.0 at large effects (Figures 6 and S1). For an allele at a starting frequency of 0.35, the design reached 80% power at a per-allele effect of 0.70 phenotypic standard deviations at 200× coverage and at 0.95 standard deviations at 100× coverage. For an allele at a starting frequency of 0.15, it reached 80% power at 0.93 and 1.28 standard deviations, respectively. These thresholds correspond to large effects, on the scale of the major color genes the scan was meant to detect. Our E&R design with two replicate populations did not reach 80% power at either starting frequency for any effect size.

The difference between the designs was swamped by background noise. At an intermediate starting frequency of 0.25, σ(p_0_) was 0.23 in our E&R data at two replicate populations and 0.05 for a case/control pool sequenced at 200× coverage (Figure S1). A case/control pool at 200× carried lower noise than the E&R design at twenty replicate populations (Figure S1).

Higher replication raised the power of the E&R design. With five replicate populations, it reached 80% power at per-allele effects of 0.15 to 0.25 standard deviations for alleles at starting frequencies of 0.05 and 0.15, but did not reach 80% power for a common allele at a starting frequency of 0.35 in any comparison. With ten replicate populations, it reached 80% power at effects of 0.12 to 0.20 standard deviations at all three starting frequencies, including the common allele. Taken together, the more powerful design depended on replication. At two replicate populations, E&R never reached 80% power at starting frequencies of 0.15 or 0.35, whereas a single generation of case/control sampling did. At five, E&R outperformed case/control for rare alleles, but only case/control reached 80% power for a common allele. E&R required ten replicate populations to reach 80% power at all three starting frequencies.

### Inversion frequencies differ among lines in one population

Cosmopolitan inversions were polymorphic only in Miami (Figure 7). Selection separated the light and dark lines. In(2L)t and In(3R)K rose in the light line (In(2L)t: control vs. light OR = 0.38, p = 6.1 × 10⁻³³; In(3R)K: control vs. light OR = 0.49, p = 3.9 × 10⁻¹⁰) but were lost in the dark line (In(2L)t: control vs. dark OR = inf, p = 7.9 × 10⁻³⁰; In(3R)K: control vs. dark OR = 97.2, p = 3.3 × 10⁻²⁹). In(3L)P was lost in both selected lines (control vs. light OR = 89.9, p = 5.9 × 10⁻²⁶⁴; control vs. dark OR = 89.6, p = 5.9 × 10⁻²⁶⁴). In(3R)Payne did not change (control vs. light OR = 0.74, p = 0.10; control vs. dark OR = 1.32, p = 0.42). The other D. melanogaster population (Toronto) and both *D. simulans* populations did not carry these inversions, so the pattern could not be tested again.

**Figure 7:**
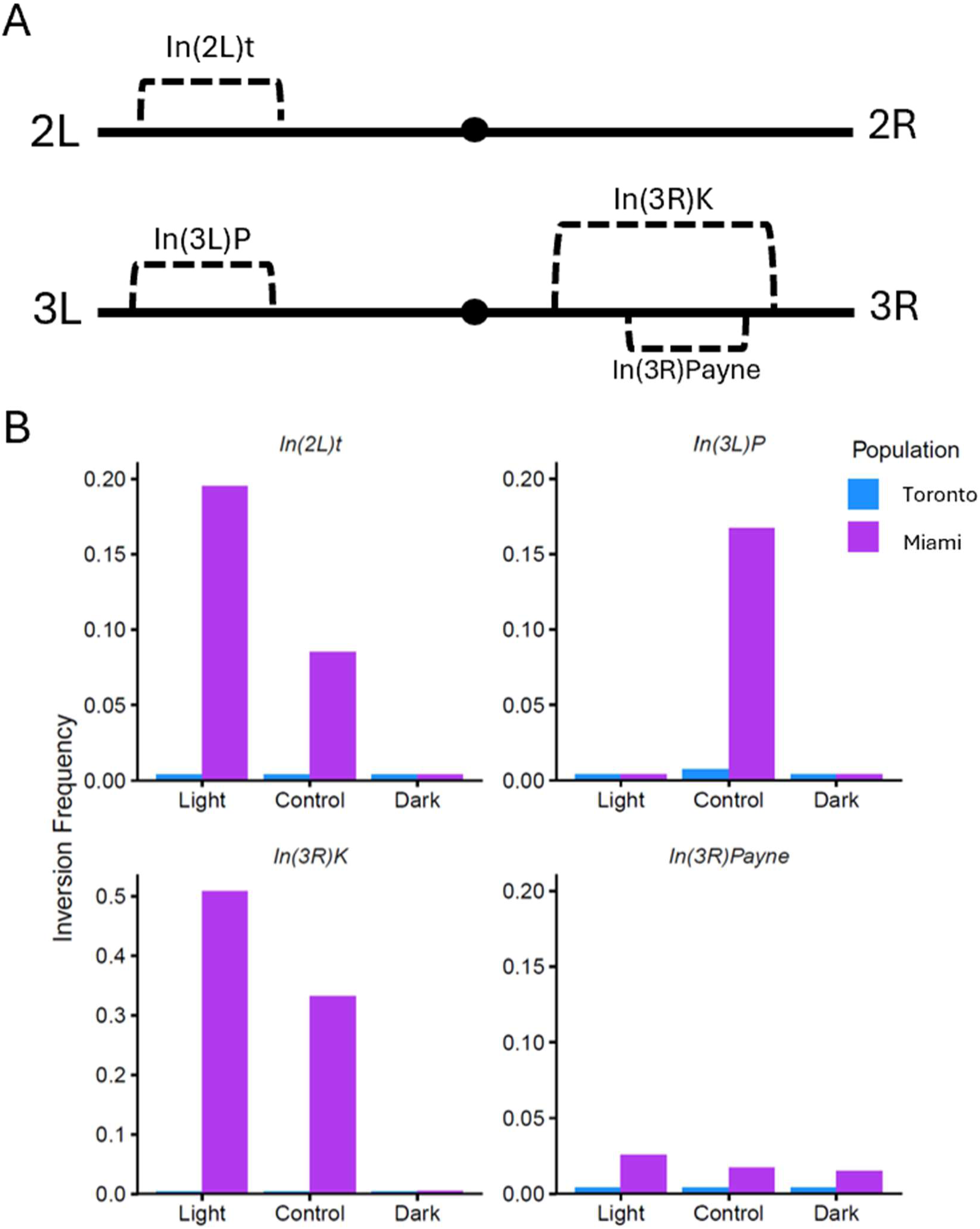
Cosmopolitan inversions decline under dark selection in the one population that carries them. Frequencies of four cosmopolitan inversions estimated from diagnostic marker SNPs, by line, population, and inversion. Inversions were polymorphic only in Miami, where In(2L)t, In(3L)P, and In(3R)K declined toward loss in the dark selected line. Toronto and both *D. simulans* populations do not carry these inversions.

### Inversions do not inflate the background noise

Cosmopolitan inversions suppress recombination across several megabases, which can cause linked variants to change frequency together and inflate the variance of allele-frequency change across the inverted region (Tobler et al. 2014; Barghi et al. 2017). We tested whether the inversions segregating in our experiment inflated this background noise. Miami carries In(2L)t, In(3L)P, and In(3R)K, whereas Toronto carries none of them, and both populations were selected in parallel under the same regime. Under the hypothesis that inversions inflate noise, the standard deviation of allele-frequency change, σ(p0), should be higher in Miami than in Toronto.

σ(p_0_) was instead lower in Miami than in Toronto in every comparison, where Δσ = σ(Miami) − σ(Toronto). In the Dark vs. Control contrast, the difference was significant at a starting frequency of 0.15 (Δσ = −0.044, 90% CI [−0.059, −0.028], p < 0.001) but not at 0.35 (Δσ = −0.003, 90% CI [−0.025, 0.012], p = 0.70). In the Light vs. Control contrast, the difference was significant at both starting frequencies (0.15: Δσ = −0.090, 90% CI [−0.109, −0.069], p < 0.001; 0.35: Δσ = −0.083, 90% CI [−0.101, −0.053], p < 0.001). The inversion-carrying population carried less background noise than the inversion-free population, not more (Table S1).

Within populations, the arms carrying inversions in Miami (2L, 3L, and 3R) were not noisier than the non-inverted autosome (2R), where Δ = σ(arm) − σ(2R). In the Dark vs. Control contrast at a starting frequency of 0.15, σ on 3L was lower than on 2R (Δ = −0.051, 90% CI [−0.073, −0.032], p < 0.001), while 3R and 2L did not differ significantly from 2R (3R: Δ = −0.021, 90% CI [−0.049, 0.003], p = 0.16; 2L: Δ = 0.029, 90% CI [−0.001, 0.058], p = 0.11). Toronto, which carries no inversion on any arm, showed the same pattern in the same contrast and frequency (3L: Δ = −0.025, 90% CI [−0.044, −0.006], p = 0.02; 3R: Δ = −0.060, 90% CI [−0.076, −0.041], p < 0.001; 2L: Δ = 0.019, 90% CI [−0.011, 0.044], p = 0.35). Across both contrasts and both starting frequencies, no inverted arm in Miami was significantly noisier than 2R except 2L in the Dark vs. Control contrast at a starting frequency of 0.35 (Δ = 0.051, 90% CI [0.024, 0.077], p = 0.01), and Toronto showed a comparable 2L excess in that comparison despite carrying no inversion (Δ = 0.025, 90% CI [0.002, 0.046], p = 0.08). The full set of arm comparisons is given in Table S2. The arm-to-arm differences therefore appear regardless of inversion content.

### Population differentiation is higher between the D. melanogaster populations

We measured genome-wide differentiation between the two control lines within each species using Hudson’s Fst in 10 kb windows (Figure 8). Differentiation was higher between the two *D. melanogaster* populations (Toronto and Miami) than between the two *D. simulans* populations (Tallahassee-1 and Tallahassee-2). Mean windowed Fst was 0.32 in *D. melanogaster* (median 0.30, n = 12,563 windows) and 0.22 in *D. simulans* (median 0.20, n = 11,622 windows).

**Figure 8:**
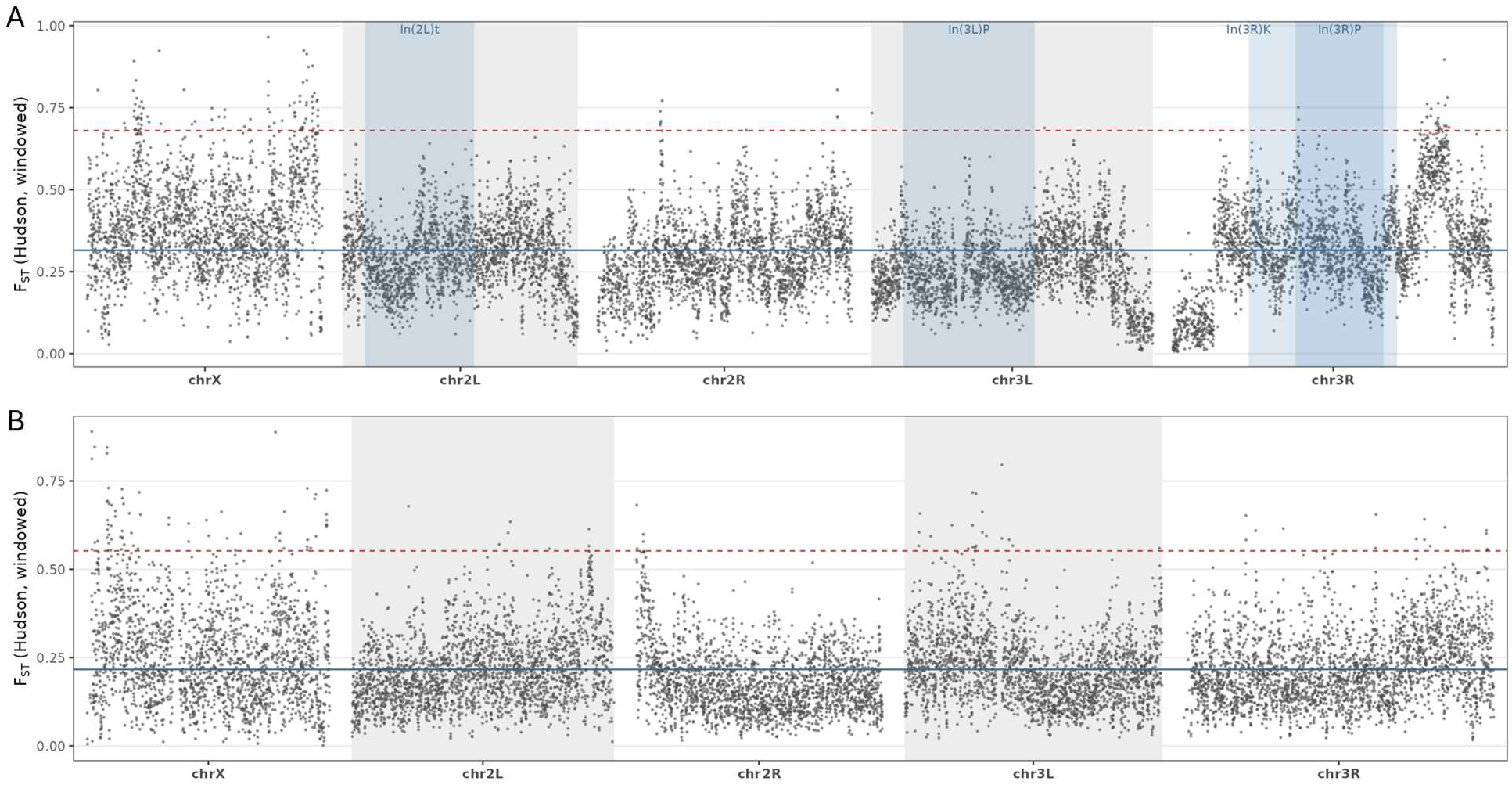
Differentiation is higher between the *D. melanogaster* populations than between the *D. simulans* samples. Hudson’s Fst in 10-kb windows along the genome between the two control lines within each species. A) Toronto vs. Miami (*D. melanogaster*). B) Tallahassee-1 vs. Tallahassee-2 (*D. simulans*). Chorion regions are excluded. Mean windowed Fst (solid blue line) is 0.32 for *D. melanogaster* and 0.22 for *D. simulans*, consistent with the two *D. melanogaster* populations being distinct localities and the two *D. simulans* samples being one population. The dashed red line indicates the windows with Fst values above 1%. The blue shaded regions are the cosmopolitan inversions (*D. melanogaster* only).

In both species, the most differentiated windows fell disproportionately on the X chromosome. Of the top 1% of windows by Fst, 88 of 126 were on the X in *D. melanogaster* and 70 of 117 in *D. simulans* (Table S3). In *D. melanogaster*, only 2 of the 126 top windows fell within a cosmopolitan inversion (In(3R)Payne), so the elevated differentiation was not concentrated over the inversions (Figure 8, Table S3).

This species difference is expected given the sampling. Toronto and Miami lie ∼2,000 km (∼1,240 miles) and 18 degrees of latitude apart, whereas Tallahassee-1 and Tallahassee-2 were collected at the same site two weeks apart. The differentiation between the two *D. simulans* samples was nonetheless substantial for populations collected from a single location. Since we measured Fst between control lines at generation 16, it includes drift that accumulated during 16 generations of selection, and much of the differentiation between the *D. simulans* samples likely reflects this drift rather than differences present at collection. The concentration of the most differentiated windows on the X chromosome, which has a smaller effective population size than the autosomes, is consistent with drift.

## Discussion

Cuticle color offers a way to separate power from architecture. Color has known genes of large effect such as *ebony* (David et al. 1985; Takahashi 2013), and it responded rapidly to selection in two species (Ruckman et al. 2026). A scan with adequate power should therefore have recovered these genes, if their large-effect alleles segregated in our populations. Instead, no individual locus emerged above the genomic background. The response to selection was nonetheless real, exceeding the differentiation expected under drift alone, which indicates that selection shifted allele frequencies across the genome rather than at a few mappable loci. A trait we expected to localize cleanly instead produced a diffuse, polygenic-looking signal in both species.

The absence of a peak is not evidence for a polygenic architecture of color. It is a property of the experiment. Modeling the power of our design against the noise in our own data shows that we had almost no power to detect a major gene of any effect size. So we could not have recovered *ebony*, or any other large-effect gene, even if it had contributed to the response in every population. This runs against the usual intuition for E&R. Selection acts in every generation, so the change in allele frequency at a selected locus accumulates, and E&R is assumed to be more powerful than a single-generation association study. However, the variance in allele frequency caused by drift grows with each generation. In our model, power in the E&R design rose with effect size but plateaued quickly, so larger effects added little power. In contrast, power in the single-generation case/control design rose with effect size without plateauing. At our level of replication, a single generation of case/control sampling would therefore have been more powerful at detecting major genes than our sixteen generations of selection. In fact, that exact design recovered *ebony* and *tan* for this trait (Bastide et al. 2013; Endler et al. 2018) in a *D. melanogaster* population from a crossed population collected in Vienna, Austria, and Bolzano, Italy. We conclude that a diffuse scan can look polygenic when it is merely underpowered, and an E&R can seem more powerful than a single-generation design when, at low replication, the reverse is actually true. The advantage of E&R over case/control is real, but it is highly contingent on replicating the E&R experiment several times from the same base population.

In addition to the detection limitation, resolution was also an issue. The ability of an E&R study to localize selected sites depends on a small number of design features, including the number of replicate populations, the duration of the experiment, the effective population size, and the level of linkage disequilibrium in the founding population (Baldwin-Brown et al. 2014; Kofler and Schlötterer 2014). Resolution improves as recombination accumulates over generations and as allele frequency changes become repeatable across replicates, both of which distinguish true targets from the neutral variants linked to them. Our design was modest on each of these axes, with two replicate populations per species, 16 generations of selection, and an effective population size that declined to 42. The founding populations were recently derived from the wild and retained substantial long-range linkage disequilibrium. We used a gradual selection regime to mitigate this by increasing selection intensity from 80% to 8.4% over 16 generations (Kofler and Schlötterer 2014; Vlachos and Kofler 2019), thereby limiting early bottlenecks and allowing recombination to break down transient linkage. However, the low effective population size, particularly in the final generations, amplified genetic drift and the stochastic decay of linkage disequilibrium, offsetting much of this benefit. The experiment was designed to drive a phenotypic response and to compare two species, not to fine-map, and the resulting resolution ceiling is shared by most E&R studies of comparable scale.

Where copy number varies, the test statistic itself becomes unreliable. Read depth at a chorion-gene cluster on chromosome 3L is elevated four- to five-fold relative to the genomic average. Under the null of no differentiation, the statistic for a 2×2 allele-count comparison scales with total coverage; as a result, the sampling variation on -log10(*p*) of a region increases in proportion to its depth. A region sequenced at five times the average therefore returns a peak roughly five times taller for the same underlying difference (since peaks are driven by outliers), and even noise-level differences can exceed genome-wide significance. Chromosome 3L illustrates this clearly. The p-value is extreme, yet the estimated allele frequency difference is large in some lines and reversed in others, a pattern that selection on a shared variant cannot produce. The chorion cluster on the X chromosome shows a similar, yet more diffuse signal. The elevation in both clusters instead reflects the developmental amplification of chorion genes in ovarian follicle cells of females during oogenesis (Spradling and Mahowald 1980), rather than a response to selection. We therefore evaluated this region by the difference in allele frequency rather than the p-value. We found no credible signal and censored it.

To our knowledge, this artifact has not been reported elsewhere, even though pooling adult females is standard practice in E&R and other pool-seq studies. Any such study should show it. Three things likely explain the omission. First, the elevation is diluted. The chorion clusters amplify roughly 60-fold on chromosome 3L and 15-fold on the X chromosome within follicle cells (Spradling and Mahowald 1980; Claycomb and Orr-Weaver 2005). However, follicle cells are a minority of the cells in a whole fly, so the signal in whole female DNA samples is only four- to five-fold higher. An elevation of this size is easily dismissed as a repetitive region or an unremarkable copy-number variant. Second, coverage is rarely inspected alongside the test statistic, and an inflated p-value is indistinguishable from a real one unless the depth is checked. Third, a spurious peak does not look anomalous in a scan that is already diffuse. In a genome-wide landscape with many weak peaks, one more peak invites a biological story rather than a technical audit.

However, limited resolution is not the only explanation for a diffuse, non-parallel response. Genetic redundancy or polygenicity could produce the same pattern, with many combinations of loci capable of generating the same color shift, and each population reaching the phenotype through a different set of variants (Barghi et al. 2019; Barghi and Schlötterer 2020). We cannot fully distinguish redundancy from low resolution, because both blur the mapping signal in similar ways. The known genetic basis of color, however, weighs against redundancy as the primary cause. If a large-effect gene such as *ebony* contributed in every population, an adequately powered scan should have detected it, and its absence is more consistent with limited resolution than with redundancy among large-effect genes.

The clearest treatment signal in our data comes from the inversions. Cosmopolitan inversions were polymorphic in only one *D. melanogaster* population, Miami. There, In(2L)t and In(3R)K rose in the light selected line and were lost in the dark, while In(3L)P was lost in both selected lines. The size and significance of these shifts argue against drift as the cause. The inversions may carry alleles for lighter color, or they may be linked to color loci elsewhere on the same chromosome arm. Endler et al. (2018) associated another cosmopolitan inversion, In(3R)Payne, with trident color in natural populations, which shows that inversions can carry trident-color alleles and makes the color interpretation plausible here, although the inversions that shifted in our experiment are not the one they implicated. We cannot distinguish these possibilities, and we cannot retest the result, because Toronto does not segregate for these inversions and *D. simulans* does not carry them. Only one population carrying the inversions is itself expected. Cosmopolitan inversions form stable latitudinal clines in North America, at higher frequency toward the equator (Kapun et al. 2016), so low-latitude Miami carries them, and high-latitude Toronto does not. Our two *D. melanogaster* populations therefore reproduce the cline and give us an inversion-carrying and an inversion-free population selected in parallel. Inversions are often assumed to degrade an E&R scan, since each suppresses recombination across several megabases, and this is a standard reason for preferring *D. simulans*, which carries few cosmopolitan inversions (Tobler et al. 2014; Barghi et al. 2017). Our contrast argues against that assumption directly. If inversions degraded E&R scans as often assumed, Miami should have been the noisier of the two populations, not the quieter one, and the inverted arms should have carried more noise than the non-inverted autosome. Neither held.

The value of color as a study system lies in this comparison. For traits such as aging and lifespan, a diffuse E&R signal is routinely interpreted as evidence of a highly polygenic architecture, and this interpretation cannot be checked, because such traits may indeed be polygenic. Human GWAS faces the same limitation, with many complex traits described as highly polygenic or even omnigenic (Boyle et al. 2017) in the absence of any way to separate genuine architecture from the limits of detection. Color removes this ambiguity. Its major genes are known, yet our scan failed to recover them. A diffuse result from a short, low-replicate experiment therefore reflects the resolution of the design at least as much as the genetic architecture of the trait. A diffuse signal is not, by itself, evidence of a polygenic trait. Long et al. (2026) demonstrated this in *Drosophila*. In a pool-seq mapping study of zinc resistance, a SNP-based GWAS found no major-effect loci, even at sequencing coverage above 700×, which is far higher than most pool-seq studies achieve. On its own, this null result would be interpreted as evidence of a polygenic architecture. Since the founders were known, the authors could estimate allele and haplotype frequencies by imputation rather than by counting alleles directly at each SNP. Imputation is far more accurate, because the error of a directly counted SNP frequency is inversely proportional to coverage and remains large even at high coverage. The imputation-based analyses recovered discrete regions of major effect in the same data. The apparent polygenicity was therefore an artifact of limited power, not a property of the trait.

Our diffuse signal contrasts with studies that mapped color successfully using different experimental designs. The most direct comparison is Endler et al. (2018), who mapped the same trait, thoracic trident color, in *D. melanogaster*. Rather than selecting over generations, they applied a pooled genome-wide association design to natural variation in two populations. They localized the trait to *ebony*, *tan*, and the cosmopolitan inversion In(3R)Payne, the very loci our scan was meant to recover. We selected directly on the trident and detected none. The same contrast holds for a related trait. Bastide et al. (2013) mapped natural variation in female abdominal color with the same pooled approach, phenotyping roughly 8,000 wild-caught females, sequencing replicate pools of the 1,000 most extreme individuals, and localizing the trait to narrow regions at *tan* and *bric-à-brac*. Both designs drew on natural populations with short-range linkage disequilibrium and large samples, which together supply the recombinant haplotypes that fine mapping requires. Their success on the same and on a related color trait indicates that our diffuse signal reflects the resolution of our experimental design rather than an intrinsic feature of color.

Pool-seq can resolve individual haplotypes when the founders are known, but our wild-derived populations provided no such reference. We also sampled a single generation and inferred inversion frequencies from tagged SNPs rather than by direct karyotyping. Sequencing males, which lack ovarian follicle cells and therefore should not show the chromosome 3L elevation, would confirm that the signal is female-limited chorion amplification. Cytological confirmation of inversion frequencies and a higher-replicate, haplotype-resolved design with known founders would directly address the remaining limitations. More broadly, resolving the genes underlying color will require a different experimental design rather than a different statistical test. Increasing the number of replicates, generations, effective population size, and the genetic diversity of the founding population would each improve resolution.

## Data Availability

Supplemental material is available at G3 online. Supplementary Figure 1 and Supplementary Tables 1–3 contain the background-noise estimates and the most differentiated Fst windows. Raw sequencing reads will be deposited at the NCBI Sequence Read Archive prior to publication. All code used for downsampling, association testing, simulation, power modeling, inversion frequency estimation, Fst, and figure generation is available at https://github.com/sruckman/Color_GWAS under the MIT license. Read alignment and allele counting used the XQTL2 pipeline (https://github.com/tdlong/XQTL2).

## Acknowledgements

We thank Matt Liu and Diego Zorio for DNA and library preparation. We used Claude Opus 5 (Anthropic) to help generate analysis code and edit the manuscript text. The authors directed its use and reviewed, verified, and take full responsibility for all code and text.

## Funding

This work was supported by National Institutes of Health awards R01OD034064 and R21AG086734.

